# The role of copper resistance in Mycobacterium tuberculosis pathogenesis

**DOI:** 10.1101/2021.03.11.434961

**Authors:** Jon Mitchell Ambler, Matthys Gerhardus Potgieter, Marisa Klopper, Melanie Grobbelaar, Margaretha De Vos, Samantha Sampson, Rob Warren, Jonathan Blackburn, Nicola Mulder

## Abstract

Despite the development of new drugs and social interventions, tuberculosis remains a leading cause of mortality. This burden falls disproportionately on developing countries, particularly those where the incidence of HIV is high. In the Western Cape, South Africa, we have identified and isolated two Beijing family strains of *Mycobacterium tuberculosis* that, despite few differences at a genomic level, differ greatly in their severity of disease caused, providing an opportunity to study virulence in this organism. The aim of this study was to identify differences at a genomic and transcriptomic level that may identify the cause of the different virulence levels observed in the two isolates.

The isolates were compared at the transcriptome level under four different growth conditions including oxidative stress. In comparing the transcriptome of the two isolates, an operon containing genes involved in the production of molybdenum cofactor that showed consistently lower levels of expression in the hypervirulent isolate was identified. A copper sensing transcriptional regulator was identified as the most probable regulator, and we found that the *Cso* operon which it is known to regulate was similarly differentially expressed in the strains.

The production of molybdenum cofactor is effected in two ways by copper levels. Through the independent insertion of copper into molybdopterin (MPT), and destabilisation of Fe-S clusters. As MoaA3 contains a Fe-S cluster that is known to be destabilised by copper, and a number of copper sensitive genes are likewise found differentially expressed, it is likely that the strains differ in terms of their levels of resistance to copper.

It is therefore hypothesised that the differences in virulence are as a result of different levels of resistance to phagosome copper overload, and the mechanism by which copper levels are linked to the production of molybdenum cofactor is described.

**Author summary:** In this article, we describe the differences in gene expression of two closely related strains of *Mycobacterium tuberculosis* isolated in the Western Cape of South Africa that differ in the severity of disease that they cause. We compared the strains at a genomic and transcriptomic level, and in doing so, we discovered a set of molybdenum cofactor genes regulated by a copper sensing transcription factor that came up in all datasets. Further genes linked to copper response were identified, providing greater evidence that the difference between the two strains was the manner in which they responded to copper stress. Phagocytes are known to exploit high levels of copper to kill intracellular bacteria, suggesting an important link between copper and disease. We conclude that resistance to copper toxicity is the most probable reason for the relative increase in virulence, and describe the regulatory relationship between copper levels and molybdenum cofactor synthesis.

## Introduction

*Mycobacterium tuberculosis* (MTB) is the causal agent of tuberculosis (TB), a chronic infectious disease that primarily infects the lungs, and can also disseminate to other organs including the lymph nodes and bones.

Despite a number of improvements in detection and treatment, the disease burden of this pathogen remains high. TB is particularly burdensome in countries with a high prevalence of HIV/AIDS (including Africa and Asia) as co-infection has a potentiating effect [1]. Those infected with HIV have a significantly greater chance of contracting TB, or for the activation of latent TB, and as both affect the host immune system treatment becomes difficult, particularly in countries with overburdened healthcare systems. According to the 2019 World Health Organisation (WHO) report, the incidence of TB in South Africa is at around 520 people per 100,000 [2], and while this number is declining, the rise of drug resistant TB threatens to rescind this progress. Of particular concern is the incidence of Beijing/W strains frequently associated with drug resistance in this region [3].

The genome of W/Beijing genotype strains differs significantly to that of the H37Rv reference strains [4–6]. Large duplications have been reported in some sublineages of the W/Beijing genotype [7] resulting in two copies of over 300 genes. This has significant implications for next generation sequencing (NGS) based experiments, making the choice of reference genome important to get an accurate picture of expression.

The isolation of two genetically similar W/Beijing genotype strains in the Western Cape provided an interesting opportunity to study the systems that determine how virulent strains of *M. tuberculosis* differ from their more benign counterparts. Despite being genetically closely related, these two strains have been characterised as hypo- and hyper-virulent based on results from a murine infection model [8]. The isolates studied are closely related with few variants observed between them, none of which were found to affect known virulence drivers in MTB.

The differences between reference genomes proved critical in the interpretation of the observed expression profiles of the two strains described in this paper, particularly in relation to the affected operon and a significant regulatory element. This provided evidence for the involvement of the molybdenum cofactor biosynthesis pathway and copper sensitivity as key elements in the differences in observed virulence.

## Results and discussion

### The effects of using different reference genomes

Within the MTB complex the H37Rv genome is by far the most well annotated and complete genome. Unfortunately, it also has fewer genes than some of the other clinically significant strains (3,906 in H37Rv vs 4,075 in W-148). The two isolates that are the focus of this project are most closely related to the W-148 isolate, and as a consequence alignment of the filtered reads to the W-148 genome produced on average 0.21% more reads than mapping to the H37Rv strain.

### Genomic comparison of the isolates

As expected, there was a high degree of genomic similarity between the two isolates. We identified 120 SNPs relative to one another, 85 of which are in coding regions. Of these 24 were classified as low impact mutations (silent), 54 as of moderate effect (missense), and 7 as high effect (frameshift mutations / stop mutations). The two isolates shared 4 large deletions relative to the W-148 reference, from 421bp to 2482bp. Some notable losses include a *esxR* gene, a PE family gene, two universal stress genes, a metalloprotease, a hypoxic response gene, a major facilitator superfamily (MFS) transporter, and an alpha/beta hydrolase. As these large deletions were found in both of the strains, they were not relevant to the comparison.

### Differentially expressed genes between isolates

Of the 102 genes found differentially expressed between the isolates, many are only differentially expressed under a subset of the growth conditions, and only 4 were found differentially expressed under all conditions. When comparing gene expression between the isolates on a per condition basis, the number of differentially expressed genes ranges from 21 to 49 (Table 1). Full lists of the differentially expressed genes are found in the supplementary tables S2-5. The most notable genes were found in one of the three operons involved in molybdenum cofactor (MoCo) biosynthesis. This cluster of genes (TBPG RS03185 - TBPG RS03200) was found to have decreased expression in SAWC5527 under all conditions (Except for TBPG RS03190 in ML(T)). The other MoCo operons and genes were not observed to be differentially expressed, implying that this operon is regulated independently.

**Table 1.** Summary of the number of genes found to be differentially expressed between the two isolates under different conditions, with isolate W-148 used as the reference genome.

| Condition | Total | Up regulated in SAWC5527 | Down regulated in SAWC5527 |
| --- | --- | --- | --- |
| Early log | 30 | 23 | 7 |
| Mid log control | 21 | 16 | 5 |
| Mid log treated | 49 | 17 | 32 |
| Stationary | 38 | 37 | 1 |

Molybdenum (Mo) is an essential microelement for nearly all organisms including MTB, where it is used by Mo containing cofactors which are in turn used by enzymes including nitrate reductase, carbon monoxide dehydrogenase (CO-DH), biotin sulfoxide reductase, as well as enzymes involved in the initial step of degradation of some pyridine derivatives [9–11].

#### Structure of the Moa operons

These genes form part of a region reportedly obtained by lateral gene transfer [12]. The expansion of the MoCo genes in the members of the MTB complex is thought to be part of the transition from an environmental generalist to an obligate pathogen [13]. The organisation of the MoCo biosynthetic genes in MTB genomes differs significantly both between and within species [14] with multiple clusters of MoCo genes found within the genome. Though these genes are annotated as having the same function, they differ significantly at the sequence level as determined by pairwise sequence alignment.

The operon in question, the *moaA3-moaB3-moaC3-moaX* gene cluster (Refered to as the Moa3 operon hence-forth), is one of the aforementioned differences between the H37Rv and W-148 genomes. In the H37Rv reference genome, *moaB3* is truncated and *moaA3* is missing due to an IS*6110* -mediated deletion [15, 16] (Figure 1). Unique to the operon in question is the *moaX* gene, a fusion of *moaD* and *moaE* (two subunits of molybdopterin synthase) which have both maintained their ability to function [14]. The operon is present in this structure in *M. bovis*, BCG, *M. tuberculosis* CDC1551, W-148 as well as in isolates SAWC507 and SAWC5527 and is thought to be the ancestral form. This structure was confirmed by sequencing of the region for the two isolate genomes, and as a result we considered the variant calling with W-148 as the reference when comparing SAWC507 and SAWC5527. No polymorphisms between the two isolates are present in the region immediately adjacent to the Moa3 operon, with the first variant seen 2,877bp upstream of the *moaA3* gene.

**Fig 1.**
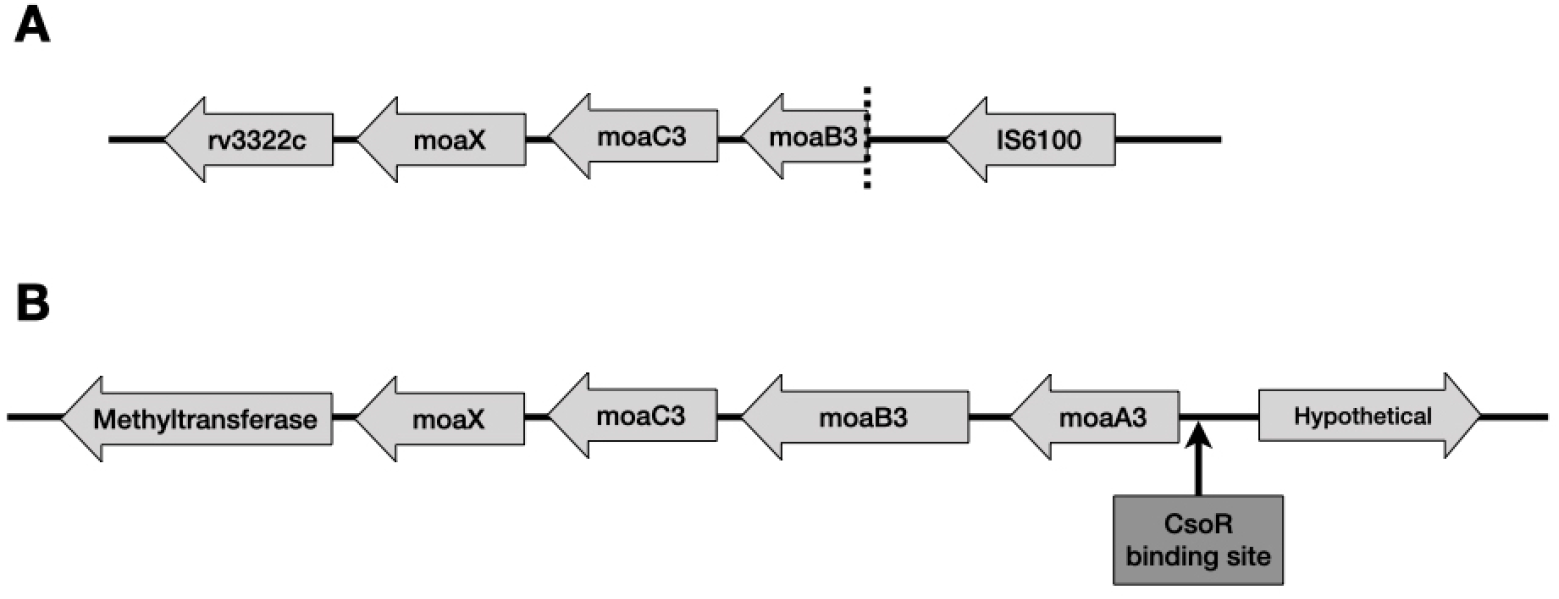
Differing structures of the Moa3 operon. **A** represents the structure in H37Rv while **B** is the structure found in isolate SAWC5537 and SAWC507.

#### Regulation of the Moa3 operon

MoaR1 is a known regulator of the Moa1 operon [17] but inspection of our WGS data showed a second rearrangement event has split *moaR1* in half in both of the isolates in question (Figure S1). This appeared to have rendered the Moa1 operon inactive, with low levels of expression under all conditions. The Moa2 operon shows some expression, but none of the genes were differentially expressed under any of the conditions tested, and is under the same transcriptional control as the Moa3 operon. Though ChIP-Seq experiments have been conducted to identify the regulators of MTB genes [18], these were conducted with H37Rv, and due to the alteration of the Moa3 operon are not applicable.

We proceeded to use *in silico* methods to identify regulators of the Moa3 operon, which led to the identification of a potential binding site 96bp upstream from the start of *moaA3* for a copper-sensitive operon repressor, CsoR (Rv0967 / TBPG RS15635). Unfortunately, *in silico* methods such as this have low specificity and the probability of a false positive is high. Further tests would be required to confirm that CsoR is a regulator of the Moa3 operon. *csoR* was observed to be significantly down regulated in the log phase in the hypervirulent SAWC5527 strain and showed lower expression in the other growth phases for SAWC5527, though not significantly. CsoR is a metalloregulatory repressor induced by copper that is known to regulate the copper sensitive operon (Cso).

This operon contains three genes (*rv0968* -*rv0970*), including *ctpV* (*rv0969*), a metal cation-transporting ATPase / efflux pump. This operon was also found to have decreased expression in isolate SAWC5527 in our datasets particularly during the early logarithmic growth phase where the expression was almost half that of SAWC507, supporting the hypothesis that it is under the same regulatory control as the Moa3 operon. Cu(I) binding to CsoR results in disassociation of the repressor from the operator-promoter region of the Cso operon [19], indicating that the lowered expression of the operon is most likely due to low levels of copper in the cell (Figure 2). The other clusters of *moa* genes did not show the CsoR binding site upstream from the TSS, indicating they are likely not under the same regulatory control by CsoR as the Moa3 operon. This is why they are not also found to be differentially expressed between the two isolates.

**Fig 2.**
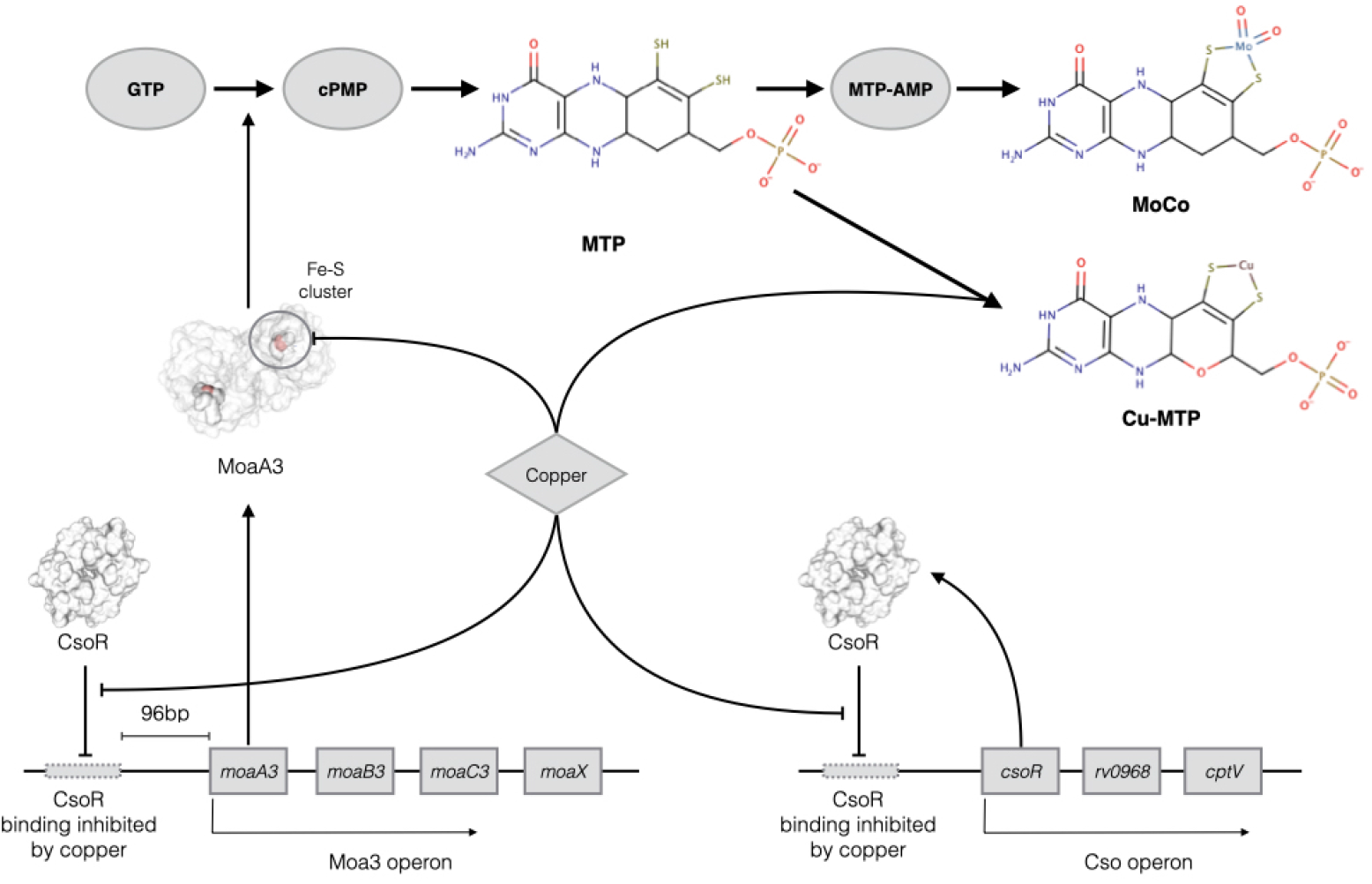
Proposed effect of copper levels on MoCo production. Under low levels of copper, CsoR is bound, lowering the expression of the Cso and Moa3 operons. Under high levels of copper CsoR binding is inhibited, allowing the expression of the Cso and Moa3 operons. In the case of the Moa3 operon this may compensate for the inhibitory action of copper on the Fe-S clusters of MoaA3, and the MoCo lost as a result of the non-specific insertion of copper into MPT. The Cso operon contains *ctpV*, which encodes a putative copper exporter that works to restore copper homeostasis.

#### The involvement of MoCo and copper resistance in pathogenesis

MoCo is an important cofactor for enzymes linked to pathogenesis in MTB [10, 13], and aids in survival in the macrophage environment. The disruption of *moaB1, moaC1, moaD1* or *moaX* has been shown to impair the ability of MTB to block phagosome maturation, reducing the ability of the bacteria to parasitise macrophages [20–22]. Some of the MoCo-dependent enzymes include the *narGHI* -encoded nitrate reductase, an important protein for MTB to survive in the oxygen deprived environment of the granuloma and associated with increasing levels of virulence in the pathogen [23, 24].

There are two notable reported mechanisms by which copper levels reduce MoCo availability. The MogA and MoeA independent insertion of copper into molybdopterin (MPT), and destabilisation of MoaA3 [4Fe-4S] clusters. Proteins including WhiB3, MoaA, and IlvD contain Fe-S clusters that are destabilised by high copper levels [25–27], and in MoaA these [4Fe-4S] clusters play a key role in the biosynthesis of molybdenum cofactor and the activity of molybdoenzymes in bacteria [28]. MoaA3 is positioned at the start of the MoCo biosynthetic pathway, where it is involved in the conversion of GTP to cPMP. Towards the end of the MoCo biosynthetic pathway, the non-specific insertion (not requiring MogA and MoeA) of copper into MPT in place of molybdenum was observed in *E. coli* and results in Cu-MPT, which acts as an inhibitor to molybdoenzymes [29]. Both of these interactions could explain why the expression of the Moa3 operon would need to be sensitive to cellular copper levels through CsoR a copper sensitive operon repressor of the Cso operon.

Metals including copper, zinc, iron and molybdenum are essential micronutrients in most forms of life, including MTB, with even small fluctuations having large effects on the ability of the pathogen to establish infection and large fluctuations being fatal to the organism. The phagosome disturbs this delicate homeostasis, attempting to overload the bacteria with toxic levels of copper [30]. In turn, MTB has been shown to respond to copper levels during infection via the aforementioned copper sensitive transcription factors [31]. Copper overload is one of the mechanisms used by macrophages to destroy MTB within their phagosomes through iron-sulfur cluster degradation [Fe-S] [25, 26, 30] and metal cofactor replacement [30].

### Other differentially expressed genes linked to copper

In our dataset, we found a number of genes linked to copper that were found to be differentially expressed between the two isolates. This included a copper sensing repressor encoding gene *ricR* (Rv0190 / TBPG RS01010), a paralogue of *csoR* previously mentioned as a ML(C) gene of interest. RicR regulates an operon containing the *lpqS* gene (Rv0847 / TBPG RS16265) which encodes a probable lipoprotein induced by copper [30, 32]. In our results, we see that *lpqS* is significantly down regulated in SAWC5527 during ML(C) and down regulated in all other conditions in SAWC5527, though below the threshold of significance. Its expression mirrored that of *mmcO*, a multicopper oxidase required for copper resistance that interacts with multiple cation transporting proteins including CtpV from the Cso operon that also showed significantly lower expression during early and middle log phase growth in isolate SAWC5527, with the expression of both genes being roughly half in this isolate. Another gene linked to *lpqS* was the metallothionein gene *mymT* (Rv4014), which was also found to be down regulated in SAWC5527 during Elog and ML(C) phase growth, and has previously been identified along with *lpqS* to be involved in the resistance mechanisms of MTB to phagosomal copper overload [30].

### Additional genes of interest

Apart from the genes with an apparent link to MoCo biosynthesis or copper metabolism, a number of differentially expressed genes were also noted as potentially contributing to the differences in phenotypes, many only observed under certain conditions. Details of the genes that were significantly differentially expressed with a-value less than 0.05 can be found in the supplementary tables S2-5.

When comparing the strains during early logarithmic phase growth, we observed genes with proximal SNPs (where isolate SAWC5527 differs from SAWC507) that may have an effect on their expression. Two examples are TBPG RS03725 and TBPG RS15510 that have SNPs 54bp, 2102bp away respectively, that could potentially be the cause of the altered expression. The first, TBPG RS03725, is an acetyl-CoA carboxylase that showed a decrease in expression in the more virulent isolate SAWC5527. The second is TBPG RS15510 (Rv0991c in H37Rv), which is annotated as a FmdB family transcriptional regulator in W-148 that has been identified as one of the genes involved in regulatory mechanisms in response to nitrogen limitation in *Mycobacterium smegmatis* [33, 34]. *This gene was also found to have decreased expression in SAWC5527 but is only 333bp long with neither of its flanking genes showing significant differential expression*.

*During middle logarithmic phase growth, we observed esxB* (TBPG RS20410 / Rv3874) showing increased levels of expression during both ML(C) and ML(T) conditions in isolate SAWC5527. This secreted virulence factor is required for pathogenesis in *Staphylococcus aureus* and MTB [35]. EsxB is secreted by the ESX-1 system along with EsxA (Rv3875) [36], which did now show any significant changes in expression in any of the observed conditions.

Treatment with *H*_2_*O*_2_ yielded the greatest number of differentially expressed genes between the two isolates, though a large number of these were annotated as hypothetical proteins. What this does reveal, is that the two strains respond quite differently to oxidative stress as induced by *H*_2_*O*_2_. Notable genes found in this dataset with links to pathogenicity were a type B diterpene cyclase (TBPG RS17815) and a diterpene synthase (TBPG RS17820) that were both found to be significantly down-regulated with half the level of expression in the hyper-virulent strain. Diterpenes have been studied in the past for their potential role in promoting phagolysosome maturation arrest [37, 38] and while it is unclear why the hyper-virulent strain would have lowered expression for these genes, it shows once again that these isolates are responding in different manners to conditions in the phagosome.

Two virulence factors shown to be involved in resistance to oxidative stress are Rv2617c (TBPG RS12610) and P36 (Exported repetitive protein Erp or Rv3810) [39]. The expression of Rv2617c (TBPG RS12610) in our dataset was shown to be significantly increased in SAWC5527 during treatment with *H*_2_*O*_2_, while P36 showed an increase but did not pass significance testing.

P36 is under the control of sigma factor H (SigH) [40] which has lower expression in SAWC5527 relative to SAWC507, though only with a significant q-value during stationary phase growth. SigH has been shown to play a crucial role in the response to environmental conditions including oxidative-stress, envelope damage, hypoxia [41, 42], survival in the phagocytes possible through the modulation of the host’s innate immune response [43], and as a key regulator of the transition phase in *Clostridium difficil* [44]. *The expression of sigH* also appeared highest during stationary phase, lower during early log phase and lowest in mid log phase growth in both samples possibly indicating a role in dormancy, and that SAWC5527 is generally in a more active state.

The stationary phase contains the largest set of adjacent differentially expressed genes (which included 7 genes from TBPG RS17790 to TBPG RS17825) found to be down-regulated in the hyper-virulent strain. These include the previously mentioned diterpene synthesis related genes found in the *H*_2_*O*_2_ treated samples. The other genes appear to be likewise involved in metabolism, including trehalose-phosphate phosphatase that is located in the cell wall and induces humoral and cellular immune responses in the host [45]. Though considered to be significantly differentially expressed, many of these genes have a very low number of mapped reads compared to other genes in the region. Specifically TBPG RS17785, TBPG RS17810, and TBPG RS17815 have a higher read coverage while the remaining genes of the cluster show low levels of coverage. Despite this, these low coverage genes had little inter-replicate variance, and are still significantly differentially expressed.

## Materials and methods

### Mycobacterial culturing and sequencing

Culturing of mycobacterial strains was conducted at Stellenbosch University, with the sequencing being done at the Agricultural Research Council in Pretoria. The goal of our study was to identify the genes involved in the altered pathogenicity of the MTB strain SAWC5527 (Hyper-virulent) compared to SAWC507 (Hypo-virulent) [8].

In order to do this, the two strains were compared at different growth phases, as well as stress conditions. Cultures were grown in Middlebrook 7H9 liquid media with the addition of dextrose and catalase, with growth phases determined by monitoring optical density over time. Samples for RNAseq were extracted during early logarithmic phase (Elog), middle logarithmic phase (ML(c)), and stationary phase (Stat). A fourth set of samples was treated during ML(C)) with hydrogen peroxide (*H*_2_*O*_2_) to simulate growth in the phagosomal environment (ML(T)). Each condition had 3 biological replicates, resulting in 24 total samples sequenced. For the removal of rRNA from the total RNA samples the Truseq stranded mRNA library preparation kit (RS-122-2101) with the Bacterial Ribozero kit (MRZMB126) was used. The samples were sequenced on an Illumina HiSeq 2500 using version 4 SBS chemistry (2×125bp), with approximately 10 million reads per sample loaded into the lane.

### Read filtering, alignment, and differential expression

Reads were trimmed and filtered using Trim Galore! (version 0.4.0, https://www.bioinformatics.babraham.ac.uk/projects/trim galore/), and aligned to both the H37Rv (Accession number NC 000962.3) and W-148 (Accession number NZ CP012090.1) isolate genomes using BWA-MEM. Alignments were filtered using SAMtools. GenGraph (https://github.com/jambler24/GenGraph) was also used to create a homology matrix to allow mapping of annotations across species. Differential expression analysis was conducted using Cuffdiff (v2.2.1) with a masking gff3 file containing rRNA and tRNA. Cuffdiff uses a Benjamini-Hochberg correction to produce a q-value, and genes with a q-value ¡ 0.05 were flagged as significantly differentially expressed.

### Variant calling

Variant calling was done using whole genome sequencing reads from SAWC5527 and SAWC507 mapped against both the W-148 and H37Rv genomes individually. The pipeline makes use of BWA-MEM for the alignment of the reads, SAMTools for the filtering of the aligned reads, and the GATK HaplotypeCaller for variant calling. The variant files were then filtered using vcftools (0.1.12b) and annotated using SnpEff [46]. To detect large chromosomal deletions, an in-house tool written in Python was used.

### In silico detection of transcription factor binding sites using FIMO

The tool Find Individual Motif Occurrences (FIMO) was used to detect transcription factor (TF) binding sites in regions that differ between the two strains [47], and a list of TF binding motifs from a genome-wide TF binding study conducted by Minch *et. al*. [48]. *In the case of the Moa3 operon, a region 200bp upstream from the start of the moaA3* gene was scanned for known motifs.

## Conclusion

Comparison of gene expression between isolates SAWC507 and SAWC5527 under different conditions showed that the genes of the Moa3 operon were consistently down regulated in the SAWC5527 isolate. In searching for regulators of this operon, we identified a disruption in *moaR1* in both the isolates, which, taken with the lack of differential expression in the Moa1 operon, appears to indicate that *moaR1* is not the driver of the altered expression seen in the Moa3 operon. Identifying the true regulator of Moa3 was made difficult by the rearrangement in the Moa3 operon found in the H37Rv reference genome. This highlighted the importance of having an accurate reference genome. By employing *in silico* methods to identify possible TF binding sites close to the Moa3 operon in the W-148 genome, which has the correct operon structure, we identified a possible CsoR TF binding site adjacent to the operon. The Cso operon was likewise found to have lowered expression, indicating that the hypervirulent SAWC5527 strain is responding as though experiencing decreased levels of intracellular copper. The cause of the different virulence phenotypes observed between the two strains may therefore be tied to their ability to resist phagosomal copper overload, a mechanism found in macrophages. While there were multiple differences between the isolates at a genomic level, none can be deemed causative, as the implicated proteins are not yet fully understood.

The most parsimonious explanation of the data is that these two strains respond differently to conditions in the macrophage (supported by the observation that the greatest number of genes were differentially expressed in the *H*_2_*O*_2_ treated samples) including the copper induced toxicity, which effects MoCo production. As the Moa1 operon is not active / responsive, it falls to the Moa3 operon to respond, potentially as a result of the regulation by CsoR. This is supported by the differential expression of a number of other copper related genes, including the Cso operon.

The mechanism of virulence described in this paper warrants further study including *in vitro* experiments, and will require the use of more closely related reference genomes, which though available, are not as well annotated.

## Supporting information

**S1 File. Site of chromosomal rearrangement in H37Rv**. This figure shows how reads from isolates SAWC507 and SAWC 5527 appear when aligned to the H37Rv reference genome in the region of the Moa3 operon where a chromosomal breakpoint is found.

**S2-S5 File. Amalgamated results of the differential expression of genes during the early log phase of growth**. These tables were generated as an output from an in-house tool named Holmes. The SNPs column refers to any variants found within the coding sequence of the gene. The operon column high-lights clusters of genes that are all differentially expressed and probably part of an operon. Any gene proximal to the differentially expressed gene that is also differentially expressed is listed. The final column shows the homologue of the gene in H37Rv as predicted by whole genome sequence alignment and annotation overlapping done by the tool GenGraph [49].

## Acknowledgments

We thank the National Research Foundation of South Africa for their financial support of this research, as well as the University of Cape Town for use of their facilities. This work was funded by the National Research Foundation of South Africa, grant number 86934. SLS is funded by the South African Research Chairs Initiative of the Department of Science and Technology and National Research Foundation (NRF) of South Africa, award number UID 86539. JMB and SLS are funded by the South African Research Chairs Initiative of the Department of Science and Technology and National Research Foundation (NRF) of South Africa, award numbers UID 64760 and 86539 respectively. The content is solely the responsibility of the authors and does not 323 necessarily represent the official views of the NRF. The authors would also like to thank 324 Dr. Monique Williams for providing input and insight for the paper before submission.

