## Supplementary figures and images for "The role of copper resistance in Mycobacterium tuberculosis pathogenesis"

### Fig-S1

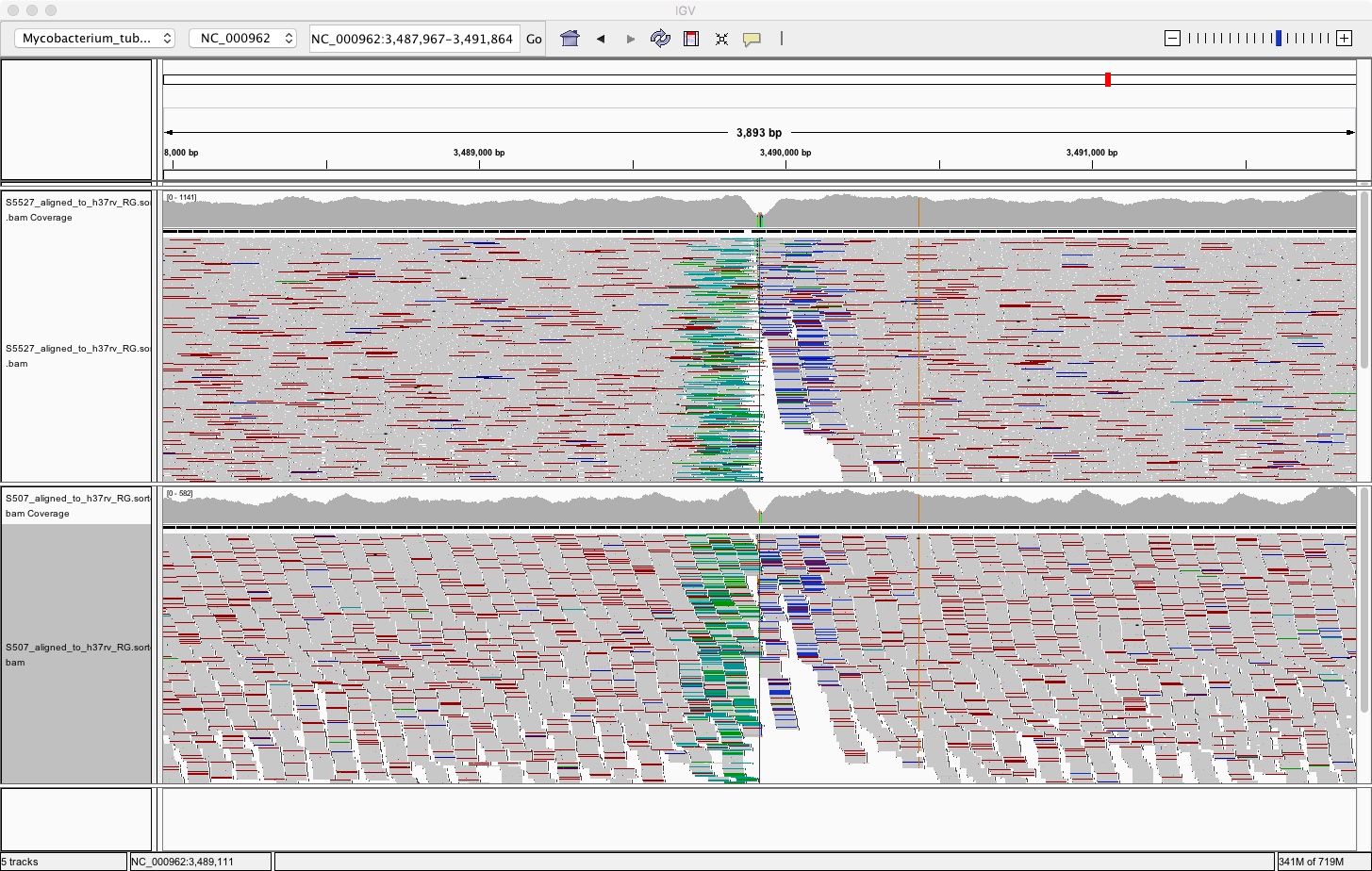
